# Non-neuronal signal fluctuations in Alzheimer’s disease and in mild cognitive impairment

**DOI:** 10.1101/2025.10.01.679773

**Authors:** Maria Guidi, Giovanni Giulietti, Taljinder Singh, Matteo Mancini, Mauro DiNuzzo, Sabrina Bonarota, Giulia Caruso, Carlotta Di Domenico, Fabrizio Esposito, Laura Serra, Carlo Caltagirone, Giovanni Augusto Carlesimo, Federico Giove

**Author notes:** Corresponding author: Federico Giove, Ph.D., Neuroimaging Laboratory, Fondazione Santa Lucia IRCCS, Via Ardeatina 306, 00179 Rome, Italy.

## Abstract

Blood oxygenation level dependent (BOLD) functional magnetic resonance imaging (fMRI) permits the investigation neural activity thanks to the neurovascular coupling mechanism. However, neural activity accounts for only a portion of the observed BOLD signal fluctuations, as the vasculature integrates multiple physiological inputs that contribute to the response. Research focusing on isolating the vascular components of the BOLD signal revealed that markers of cerebrovascular health, such as cerebrovascular reactivity (CVR), serve as valuable biomarkers for neurodegenerative diseases.

This study examines the relationship between vascular metrics and noise in a cohort comprising individuals with Alzheimer’s disease (AD), mild cognitive impairment (MCI), and healthy controls (HC). Vascular responses were assessed using three functional contrasts during a hypercapnic challenge: arterial spin labeling (ASL) to measure cerebral blood flow (CBF) reactivity, vascular space occupancy (VASO) to quantify cerebral blood volume (CBV) reactivity, and BOLD imaging. Noise metrics were derived from multi-echo BOLD resting-state data by isolating the TE-independent components of the signal.

Mean correlation coefficients for noise vs ASL-CVR are: (–0.12 ± 0.06) for HC, (–0.14 ± 0.08) for MCI, (–0.11 ± 0.05) for AD. Mean correlation coefficients for noise vs BOLD-CVR are: (0.25 ± 0.11) for HC, (0.24 ± 0.07) for MCI, (0.23 ± 0.11) for AD. Mean correlation coefficients for noise vs VASO-CVR are: (0.13 ± 0.10) for HC, (0.13 ± 0.07) for MCI, (0.12 ± 0.12) for AD. These results suggest that TE-independent noise relates to the three vascular contrasts to varying extents and directions, with no significant differences across groups. Further analysis within specific functional networks revealed group differences in specific networks.

The observed cortical correlations between noise and vascular features provide important insights into brain function and the progression of neurodegenerative diseases, offering a potential avenue to disentangle vascular and neural contributions in brain network and connectivity studies.

## Introduction

Blood oxygen level dependent (BOLD) functional magnetic resonance imaging (fMRI) indirectly detects neural activity via the associated vascular response. The vasculature determines the observed BOLD signal mainly with two components: a static one, given by the vascular organization and blood density distribution, and a dynamic one, that depends on the active dilatory capacity of the vessels.

Neural activity is one of the many inputs to the vasculature that are capable of eliciting a vascular effect and, in turn, a detectable BOLD signal change. Other inputs include physiological processes such as respiration, heartbeat, fluctuations in partial pressure of blood CO_2_, Mayer waves, etc. (T. T. Liu, 2016; Tong, Hocke, & Frederick, 2019; Wise, Ide, Poulin, & Tracey, 2004).

Noise in fMRI includes system-related noise (of thermal and instrumental origin) and subject-related noise, due to the above mentioned non-neuronal physiological processes and motion. While in general anything non-neuronal may be treated as noise, the exact definition depends on the specific application and target.

The vascular component for example, as opposed to the neuronal component, may be considered a confound in fMRI studies, since it can give rise to amplified or delayed responses that do not reflect a different neuronal engagement or actual lagged neuronal activity, but rather different underlying vascular features (Chang, Thomason, & Glover, 2008; Mangini et al., 2025). However, the vascular response might represent the component of interest in specific applications given its informative nature on the brain vascular health status. In this regard, mapping the cerebrovascular reactivity (CVR) has proven useful in disease characterization and in aging studies (Chen, 2018; P. Liu et al., 2021; Sleight, Stringer, Marshall, Wardlaw, & Thrippleton, 2021). CVR is a measure of the ability of blood vessels to dilate or constrict upon vasoactive stimuli; in fMRI it is usually assessed by increasing the arterial partial pressure of CO_2_ while acquiring a perfusion-sensitive contrast and it is expressed in units of %/mmHg (percent signal changes over unit increase in partial pressure of end-tidal CO_2_). In Alzheimer’s disease (AD), cerebral blood flow (CBF) and CVR reduction were found to be early changes and indicative of symptom severity (Nielsen et al., 2017), thus offering a great potential for the characterization of the disease before symptom manifestation.

In this work, vascular metrics are compared to noise metrics derived from a set of fMRI acquisitions in a healthy control (HC) group, in a group of individuals with mild cognitive impairment (MCI) and in a group of individuals with AD. The vascular metrics are derived from a respiratory-challenge task executed during the consecutive acquisition of two sequences: a pulsed arterial spin labeling sequence (PASL) and a vascular space occupancy (VASO) sequence, which also provides a simultaneous BOLD timeseries. Noise maps are computed from an independent resting-state multi-echo BOLD timeseries by retaining the TE-independent components.

## Materials and Methods

The recruited group consists of 43 healthy controls (HC), 18 subjects with mild cognitive impairment (MCI), and 18 subjects with Alzheimer’s disease (AD). The mean age ± std is 65.3 ± 7.4 y for the HC group, 72.1 ± 4.2 y for the MCI group and 71.9 ± 5.5 y for the AD group. Sixteen subjects (5 from the AD group, 5 from the MCI group, and 6 from the HC group) were excluded due to insufficient data quality of the respiratory recordings.

### Data acquisition

Data acquisition was conducted at Santa Lucia Foundation IRCCS in Rome using a Siemens MAGNETOM 3T MRI scanner, in accordance with the ethical standards of the Helsinki declaration and its later amendments. The study protocol was approved by the local ethics committee (Approval 2022_010), and written informed consent was obtained from all participants prior to the study. The protocol included the following sequences for a total acquisition time (TA) of 22 min: a VASO functional sequence (Huber et al., 2023) (nominal resolution: 2.0 × 2.0 × 3.0 mm^3^, TR = 2.7 s, TE = 14.3 ms, flip angle = 30° with variable flip angle scheme, inversion delay = 550 ms, CAIPIRINHA 3×1, TA = 10 min), a PASL functional sequence (Günther, Oshio, & Feinberg, 2005) (nominal resolution: 4.4 × 4.4 × 4.4 mm^3^, TR = 5.5 s, TE = 13.3 ms, flip angle = 120°, perfusion mode FAIR QII, partial Fourier (phase and slice) = 6/8, CAIPIRINHA 2×2, TA = 10 min), a PASL sequence with reverse phase encoding (TA = 30 s), and a PASL M0 sequence (TR = 6 s, no background suppression, TA = 54 s).

The respiratory paradigm, which was repeated twice in each session (once during the VASO acquisition and once during the PASL acquisition), consisted of an alternation of medical air and gas mixture (5% CO_2_, 21% O_2_, 74% N_2_) delivery. The mixtures were switched every 2 minutes, starting with medical air and until the end of the functional acquisition. The gases were administered through a face mask; participants additionally had a nasal cannula for end-tidal CO_2_ and O_2_ recording, and a pulse plethysmograph and respiratory belt for physiological monitoring.

In another session, the participants underwent a functional BOLD acquisition in resting-state conditions (nominal resolution: 2.8 mm, TR = 0.98 s, TE1/TE2/TE3 = 12.8/30.42/48.02 ms, flip angle = 63°, multiband acceleration factor = 4, TA = 8 min). During the acquisition, participants stared at a fixation cross and were instructed not to fall asleep.

### ASL data analysis

The ASL4D data (control-label pairs) were analyzed with the goal of obtaining cerebral blood flow (CBF) time series (CBF4D) in absolute units. The processing pipeline consisted of two major stages: *(i)* ASL4D preprocessing, including motion correction, skull stripping, smoothing, computation of perfusion-weighted images (PWI4D), and denoising; and *(ii)* CBF quantification using the ExploreASL framework.

Head motion in the ASL4D data was corrected using a realignment strategy that accounted for the alternating control–label acquisition scheme. Specifically, all volumes were realigned to the mean ASL4D image using the zig-zag approach (Wang, 2012), based on six-parameter rigid-body registration as implemented in SPM12. This method first performs a conventional realignment to the mean ASL image and then regresses out a zig-zag regressor (1, −1, 1, −1, …; with ‘1’ denoting a control volume and ‘−1’ a label volume) from the motion parameters. This step removes variance specifically related to the alternating label–control sequence and helps reduce spurious motion-related artifacts in the final dataset. The corrected parameters were then applied to the original images to generate the realigned ASL4D dataset (rASL4D). The three volumes acquired for the M0 sequence were averaged to create a mean M0 image, which was subsequently coregistered to the temporal mean of the rASL4D using ANTs. The resulting M0 image (rM0) was skull-stripped to generate a binary brain mask in ASL space. This mask was later applied to remove extra-cranial signal and to ensure that subsequent analyses were restricted to brain tissue. After motion correction and masking, spatial smoothing was applied to rASL4D using an isotropic Gaussian kernel with a 5 mm full width at half maximum (FWHM). This generated the smoothed dataset (srASL4D). Perfusion-weighted images (PWI4D) were then computed as the voxelwise difference between control and label images from srASL4D. The brain mask derived from the M0 sequence was applied to these perfusion images to exclude non-brain regions. Following these initial steps, the PWI4D dataset was denoised using the patch-based Low-rank and Sparse Tensor decomposition combined with Non-Local Means filtering (pLST-NLM) algorithm (He, Lu, Li, Lu, & Zhu, 2022). This hybrid method combines two complementary strategies: first, low-rank and sparse tensor decomposition (pLST) extracts a low-rank representation of the temporal signal while suppressing sparse noise; second, non-local means (NLM) filtering reduces residual Gaussian noise by weighting voxel intensities according to patch similarity. We applied this method using a three-step sequence: first pLST, then NLM on the pLST output, followed by a final pLST step on the NLM-filtered images. The resulting denoised perfusion series served as the primary ASL input for subsequent CBF quantification.

The denoised PWI4D, together with rM0, T1-weighted (T1w), and FLAIR images, were processed with ExploreASL (Mutsaerts et al., 2020). ExploreASL includes two major modules: a structural module for anatomical processing and an ASL module for CBF quantification. The structural module began with linear registration of the T1w image to MNI space (AC–PC alignment), ensuring consistent anatomical alignment for quality control and group-level analyses. The FLAIR image was then registered to the T1w image, enabling integration of FLAIR-derived information (e.g., lesion masks) into the T1w space. White matter hyperintensities (WMH) were segmented on the FLAIR image, and lesion filling was applied to the T1w image to mitigate misclassification errors during tissue segmentation. Tissue segmentation of the T1w image was then performed, generating probability maps for gray matter (GM), white matter (WM), and cerebrospinal fluid (CSF). Whole-brain tissue volumes were also estimated from these maps. The ASL module processed the perfusion-weighted and M0 images. First, PWI4D and rM0 were registered to the T1w space. Partial volume (PV) maps were prepared by reslicing GM and WM tissue probability maps to ASL resolution and smoothing them, enabling correction for PV effects. The M0 image underwent preprocessing to remove intensity inhomogeneities and generate a bias field for normalization. An analysis mask was also created to exclude regions affected by susceptibility or vascular artifacts.

CBF quantification was then performed by combining PWI4D with the processed M0 image. The quantification step included blood T1 correction, estimation of labeling efficiency, and acquisition-specific parameters, including labeling strategy (PASL), readout type (3D GRASE), post-labeling delay (2000 ms), and vendor-specific scaling. Partial volume correction was applied using the method of Asllani et al., which accounts for tissue mixing effects at the voxel level, improving the accuracy of CBF estimates (Asllani, Borogovac, & Brown, 2008). Finally, perfusion maps quantified in absolute physiological units (ml/100 g/min) were generated and normalized to the MNI152 space (Mazziotta, Toga, Evans, Fox, & Lancaster, 1995) with a resampled resolution of 2 mm.

### VASO and BOLD data analysis

The VASO sequence used for the acquisition provides a simultaneous BOLD timeseries, since it acquires an alternation of BOLD and VASO-weighted volumes (Huber et al., 2021).

The timeseries was first corrected for motion similarly as described for the ASL preprocessing. Subsequently, the VASO-weighted and BOLD timeseries are split and upsampled in time, effectively doubling the number of data points. A specific correction algorithm for the BOLD contamination in the VASO-weighted timeseries was then applied, using both the VASO-weighted data and the upsampled BOLD data to produce a corrected VASO series (Huber et al., 2021). Both the upsampled BOLD data and the VASO data were subjected to bandpass filtering. Then, a specialized denoising was employed separately for the VASO and BOLD data. The bandpass-filtered VASO and BOLD time series were fed into the denoising pipeline along with a brainmask, which restricts denoising operations to brain voxels, preventing noise from non-brain regions from influencing the brain signal. The denoising was based on the pLST algorithm and process parameters were optimized on the VASO and BOLD contrasts specifically.

The result of this process was a denoised VASO series and a denoised BOLD series, which were then normalized to the MNI152 space (Mazziotta et al., 1995) with a resampled resolution of 2 mm.

### CVR estimation

The cerebrovascular reactivity calculation was conducted using seeVR (Bhogal, 2021). Inputs to the toolbox were either the preprocessed ASL, BOLD, or VASO timecourses and the relative end-tidal CO_2_ (EtCO_2_) recording. EtCO_2_ traces were temporally aligned to the functional acquisitions by subtracting system delays before input to the toolbox. Lag-corrected EtCO_2_ traces were then estimated via cross correlation, and contrast-specific CVR maps were generated using a regression analysis (Bhogal, 2021).

### Noise map calculation

The multi-echo resting state BOLD (rs-BOLD) timeseries was automatically preprocessed with fMRIPrep and subsequently with tedana (DuPre et al., 2021). Tedana exploits the echo time (TE) dependence of the BOLD signal components to distinguish “likely BOLD” from “unlikely BOLD” components. It does so by performing first a principal component analysis (PCA) for Gaussian noise removal and then an independent component analysis (ICA) on the optimally-combined BOLD timeseries. The components are then classified based on TE dependence: TE-dependent components are accepted and TE-independent components are rejected, thus generating a denoised timecourse. Noise timecourses were then built by subtracting the denoised timecourse from the original timecourse; the result of this subtraction is therefore a timeseries including some thermal and some “non-BOLD-like” physiological noise. The temporal standard deviation of the noise timecourses were calculated and used as noise maps for the correlation analysis.

### Correlation between CVR and noise

Voxelwise scatterplots of ASL-/BOLD-/VASO-CVR vs noise were obtained for each participant both in a GM mask and within the functional networks defined by the Yeo’s parcellation (Yeo et al., 2011). For each scatterplot a correlation coefficient was obtained and averaged within group.

## Results

The averaged temporal standard deviation of noise for the HC and the MCI+AD groups are reported in Figure 1. Noise appears to be higher in the CSF compared to other compartments in both groups.

**Figure 1.**
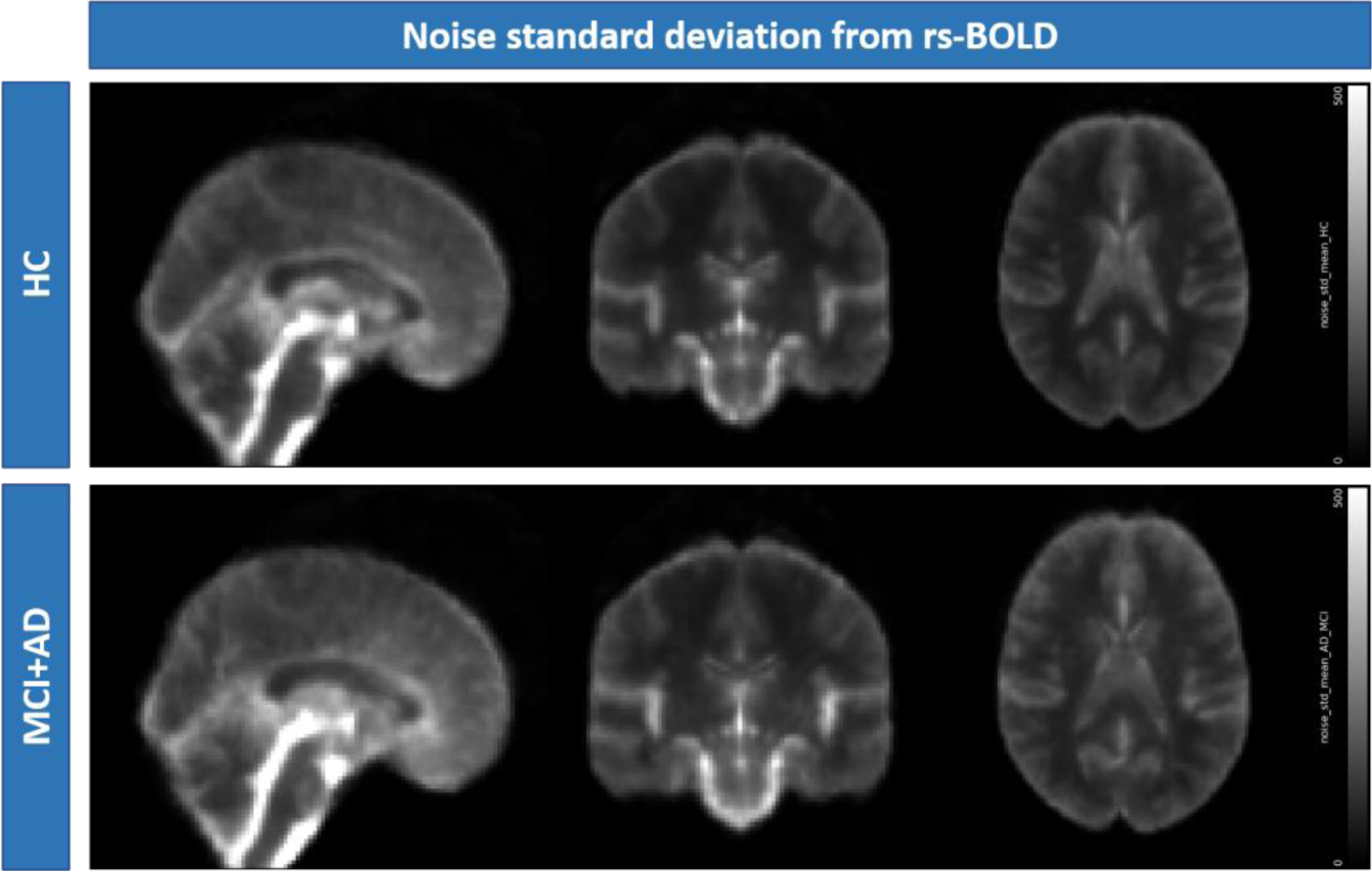
Mean maps of noise standard deviation for the HC group (top) and for the MCI+AD group (bottom). Values are in arbitrary units.

The resulting CVR maps averaged for the HC and MCI+AD groups and the corresponding mean values in GM are reported in Figure 2 and Table 1 respectively. Mean GM ASL-CVR are: (3.96 ± 1.69) %/mmHg for HC, (3.07 ± 1.80) %/mmHg for MCI, (2.68 ± 2.09) %/mmHg for AD. Mean GM BOLD-CVR are: (0.080 ± 0.023) %/mmHg for HC, (0.067 ± 0.019) %/mmHg for MCI, (0.068 ± 0.029) %/mmHg for AD. Mean GM VASO-CVR are: (0.040 ± 0.017) %/mmHg for HC, (0.034 ± 0.015) %/mmHg for MCI, (0.034 ± 0.025) %/mmHg for AD. Higher values are found for the HC group compared to the MCI and AD group, confirming a reduced vascular compliance in those groups. Significant reductions in GM were found for the AD vs HC group in ASL-CVR (p = 0.04) and for MCI vs HC group in BOLD-CVR (p = 0.02).

**Table 1.** Mean (± std) CVR values in GM for the three groups and three contrasts.

|  | ASL-CVR (%/mmHg) | BOLD-CVR (%/mmHg) | VASO-CVR (%/mmHg) |
| --- | --- | --- | --- |
| HC | 3.96 ± 1.69 | 0.080 ± 0.023 | 0.040 ± 0.017 |
| MCI | 3.07 ± 1.80 | 0.067 ± 0.019 | 0.034 ± 0.015 |
| AD | 2.68 ± 2.09 | 0.068 ± 0.029 | 0.034 ± 0.025 |

**Figure 2.**
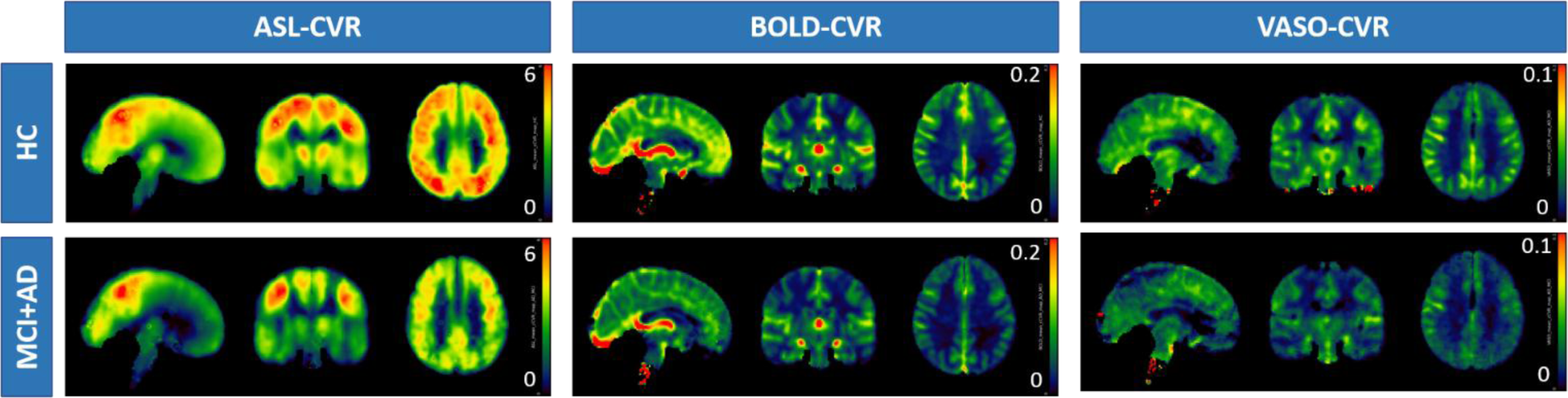
Mean ASL-CVR (left column), BOLD-CVR (central column), VASO-CVR (right column) maps for the HC group (top row) and for the MCI+AD group (bottom row). Values are in units of %/mmHg.

Scatterplots of the three CVR vs noise in GM for one representative participant are shown in Figure 3. Mean correlation coefficients for noise vs ASL-CVR are: (–0.12 ± 0.06) for HC, (–0.14 ± 0.08) for MCI, (–0.11 ± 0.05) for AD. Mean correlation coefficients for noise vs BOLD-CVR are: (0.25 ± 0.11) for HC, (0.24 ± 0.07) for MCI, (0.23 ± 0.11) for AD. Mean correlation coefficients for noise vs VASO-CVR are: (0.13 ± 0.10) for HC, (0.13 ± 0.07) for MCI, (0.12 ± 0.12) for AD. No significant group differences were found for the cortical correlation coefficients.

**Figure 3.**
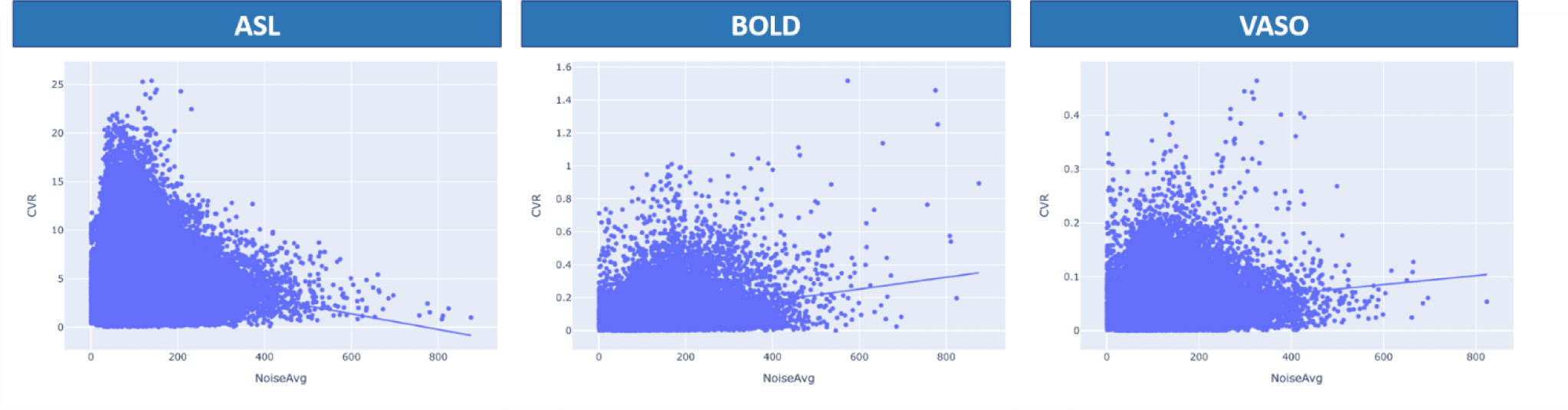
Voxelwise scatterplots of noise standard deviation vs ASL-CVR (left), BOLD-CVR (center), VASO-CVR (right) for one representative participant taken from the HC group. Voxels are taken from a GM mask.

Within-network scatterplots for one representative participant are reported in Figure 4. Mean values for the cortical and network correlation coefficients are reported in Table 2. Significant differences performing a two-tailed t-test were found for the following networks and functional contrasts: dorsal attention, HC-MCI BOLD, p = 0.01; fronto-parietal, HC-MCI BOLD, p = 0.008; fronto-parietal, HC-AD ASL, p = 0.01; default, MCI-AD ASL, p = 0.04.

**Table 2.** Mean correlation coefficients obtained from the voxelwise scatter plots of CVR vs noise for GM and for the seven networks defined by the Yeo’s functional parcellation.

| Region | Group | BOLD | ASL | VASO |
| --- | --- | --- | --- | --- |
| Cortex | HC | 0.252204706 | -0.12361971 | 0.127943218 |
|  | MCI | 0.235850232 | -0.13782446 | 0.130798542 |
|  | AD | 0.227988437 | -0.10911876 | 0.1209539 |
| Visual network | HC | 0.427668464 | -0.11038882 | 0.220527516 |
|  | MCI | 0.445452014 | -0.08313618 | 0.23560901 |
|  | AD | 0.370983718 | -0.10676654 | 0.207062722 |
| Somatomotor network | HC | 0.281718516 | -0.17248385 | 0.186179238 |
|  | MCI | 0.230896542 | -0.21437495 | 0.154956237 |
|  | AD | 0.318246151 | -0.18812584 | 0.188334188 |
| Dorsal-attention network | HC | 0.29118209 | -0.0794254 | 0.139836614 |
|  | MCI | 0.193221503 | -0.08612092 | 0.12989545 |
|  | AD | 0.238374421 | -0.02504622 | 0.117922997 |
| Ventral-attention network | HC | 0.206067706 | -0.14281389 | 0.130405651 |
|  | MCI | 0.180584272 | -0.12951908 | 0.083941288 |
|  | AD | 0.215167086 | -0.13135016 | 0.092741216 |
| Limbic network | HC | 0.090216989 | -0.05086117 | 0.070840341 |
|  | MCI | 0.060063532 | -0.02742714 | 0.057441685 |
|  | AD | 0.074400257 | 0.007367567 | -0.00604419 |
| Fronto-parietal network | HC | 0.165634232 | -0.04958759 | 0.04942645 |
|  | MCI | 0.081070398 | -0.02768995 | 0.004495187 |
|  | AD | 0.153037658 | -0.00146055 | 0.048415325 |
| Default mode network | HC | 0.18523771 | -0.10095754 | 0.083881305 |
|  | MCI | 0.160836639 | -0.12820097 | 0.060120821 |
|  | AD | 0.186757284 | -0.05799655 | 0.085123487 |

**Figure 4.**
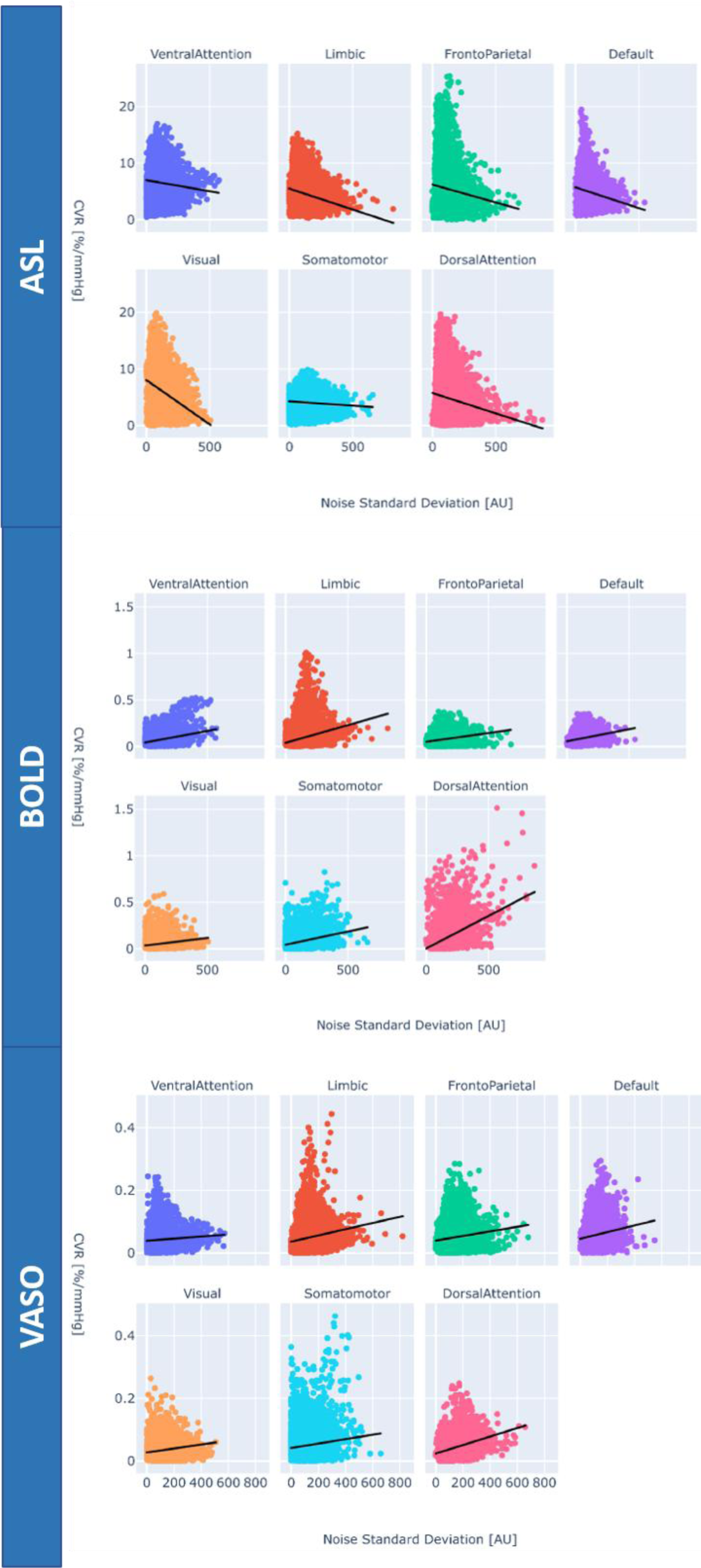
Voxelwise scatterplots of noise standard deviation vs ASL-CVR (top), BOLD-CVR (middle), VASO-CVR (bottom) for the networks defined by the Yeo’s atlas for one representative participant taken from the HC group.

## Discussion

The current work provides a link between vascular function metrics (ASL-CVR, BOLD-CVR, VASO-CVR) and TE-independent noise components extracted from multi-echo BOLD fMRI data using tedana. The observation of lower CVR measures in the MCI and AD groups compared to HC is in line with the expected vascular impairments associated with neurodegenerative disease progression (Nielsen et al., 2017). The negative correlations found between ASL-CVR and noise, coupled with positive correlations between BOLD-CVR and noise, highlight the complex vascular-neuronal interplay that contributes to functional signal variability beyond pure neural activity.

This study supports prior findings that cerebrovascular health significantly influences BOLD signal characteristics, emphasizing the importance of accounting for vascular contributions when interpreting functional connectivity and neural activation in aging and disease states (Biswal, Kannurpatti, & Rypma, 2007; DuPre et al., 2021; Kundu, Inati, Evans, Luh, & Bandettini, 2012). Using advanced multi-echo denoising approaches like tedana to isolate TE-independent physiological noise components enhances the ability to distinguish neural from non-neural signal fluctuations, reducing confounds and improving the specificity of fMRI biomarkers (Kundu et al., 2017; Lynch et al., 2020).

Furthermore, the detection of group differences within specific functional networks suggests that vascular dysfunction associated with neurodegeneration may selectively affect network-level brain dynamics, with potential implications for altered connectivity patterns observed in AD and MCI (Agosta et al., 2012; Kisler, Nelson, Montagne, & Zlokovic, 2017). These network-specific findings underscore the utility of combining vascular reactivity with noise measures to gain insights into pathophysiological mechanisms.

The present results also align with evidence showing that hemodynamic scaling and calibration of BOLD signals improve the interpretability of functional imaging data by compensating for individual differences in vascular reactivity and anatomy (Biswal et al., 2007; P. Liu et al., 2013). Hence, incorporating complementary vascular contrasts (ASL, VASO) and BOLD provides a more comprehensive characterization of cerebrovascular status especially in conditions where the neurovascular coupling might be disrupted.

Future work could extend these findings by longitudinally tracking vascular and noise metrics to better delineate the temporal trajectory of neurovascular decline and its relation to cognitive deterioration. Moreover, integrating multi-modal imaging and computational modeling approaches may further unravel the complex interactions between vascular and neuronal components, ultimately enhancing the fidelity of fMRI biomarkers for neurodegenerative disease characterization.

## Acknowledgments

The authors thank Edoardo D’Andrea for support with data preprocessing, Elena Russo for critical manuscript reading, Claudia Marzi and Marco Clemenzi for radiographical assistance, Steve Gazzitano for participant recruitment and maintenance of the gas delivery system, Alex Bhogal for helpful discussion and support with the seeVR toolbox.

## Author Contributions Statement

FG, LS, CC, GAC conceptualized and designed research. MG, FG, TS performed experiments. SB, GC, CDD recruited participants. MG, GG, TS, MM, MDN analyzed data. MG, GG wrote the main manuscript text. MG, GG, MM prepared figures. All authors reviewed and approved the manuscript.

## Funding

Work supported and funded by: #NEXTGENERATIONEU (NGEU); the Ministry of University and Research (MUR); the National Recovery and Resilience Plan (NRRP); project MNESYS (PE0000006, to NT) – A Multiscale integrated approach to the study of the nervous system in health and disease (DN. 1553 11.10.2022). MG was supported by the Italian Ministry of University and Research (MUR) - PNRR - NextGenerationEU, Mission 4 'Education and Research' - Component 2 'From Research to Enterprise' Investment 1.2 'Funding of projects submitted by young researchers' – CUP F87G25000060006.

## Disclosure/Conflict of interests

The authors declare no conflict of interest.

## Competing financial interests

The authors declare no competing financial interests.

## Notes

### Competing Interest Statement

The authors have declared no competing interest.

### Summary of Updates

Funding information was incomplete and has been modified to cite the relevant award for MG.

